# LongPhase: an ultra-fast chromosome-scale phasing algorithm for small and large variants

**DOI:** 10.1101/2021.09.09.459623

**Authors:** Jyun-Hong Lin, Liang-Chi Chen, Shu-Qi Yu, Yao-Ting Huang

## Abstract

Long-read phasing has been used for reconstructing diploid genomes, improving variant calling, and resolving microbial strains in metagenomics. However, the phasing blocks of existing methods are broken by large Structural Variations (SVs), and the efficiency is unsatisfactory for population-scale phasing. This paper presents an ultra-fast algorithm, LongPhase, which can simultaneously phase single nucleotide polymorphisms (SNPs) and SVs of a human genome in ∼10-20 minutes, 10x faster than the state-of-the-art WhatsHap and Margin. In particular, LongPhase produces much larger phased blocks at almost chromosome level with only long reads (N50=26Mbp). We demonstrate that LongPhase combined with Nanopore is a cost-effective approach for providing chromosome-scale phasing without the need for additional trios, chromosome-conformation, and single-cell strand-seq data.

## Background

Accurate reconstruction of diploid genomes is essential for a variety of studies including imprinting, disease association, and population genetics. Most genome assembly algorithms collapse the diploid into a single reference with the variants encoded at each heterozygous locus. Hence, the reconstruction of the paternal and maternal haplotypes, called phasing, is necessary for diploid genome sequencing. When sequenced by short reads, phasing relies on the extent of linkage disequilibrium in populations or the Mendelian consistency in pedigrees (e.g., the HapMap and 1000 genome projects) [1]. In recent years, long-read sequencing, e.g., PacBio and Nanopore, is now routinely used for sequencing and assembling human genomes. Consequently, phasing based solely on long reads has become the preferred methodology, because of no biases against novel variations or recombination absent in the population/pedigree.

Although there are numerous methodologies of read-based phasing developed in the last decade, only a few have demonstrated their applicability for long-read sequencing. WhatsHap is one of the most popular tools used in a wide range of applications [2, 3]. It classifies long reads into two groups by solving an NP-hard problem called minimum error correction. However, although reads are down-sampled to fifteen folds by default, WhatsHap still requires quite a few hours for phasing a human genome. On the other hand, HapCut2 is capable of assembling haplotypes using PacBio with additional proximity-ligation sequencing [4]. However, the method is still lacking validations on the error-prone Nanopore sequencing. Margin, previously named MarginPhase, constructs a hidden Markov model for simultaneous genotype calling and haplotype phasing [5, 6]. It not only reduces the running time of phasing a human genome to 1-2 hours, but also maintains high accuracy and contiguity on both PacBio and Nanopore platforms.

Although WhatsHap and Margin have proved excellent phasing accuracy on the PacBio and Nanopore platforms, their computational efficiency, taking hours for a human genome, is still unsatisfactory for large-scale phasing projects. Furthermore, because these algorithms focus on phasing single nucleotide polymorphisms (SNPs), their phasing blocks are often broken by large structural variations (SVs) (e.g., inversions and insertions) (Figure 1). It is mainly owing to the fragmentation of long-read alignments across SVs. Because phased SVs are also highly demanded in understanding the structural evolution of cancer genomes (e.g., chromothripsis) [7], a number of approaches have been developed [8, 7, 9, 10, 11]. However, these methods rely on first partitioning reads into paternal and maternal groups via SNP phasing. Subsequently, the reads in each group are used for inferring the haplotype origin of each SV. These two-stage methods are not only time-consuming, but the sizes of haplotype blocks are still bounded by the initial SNP phasing. Hence, the phasing range of existing algorithms still fail to reach the chromosome level.

**Figure 1.**
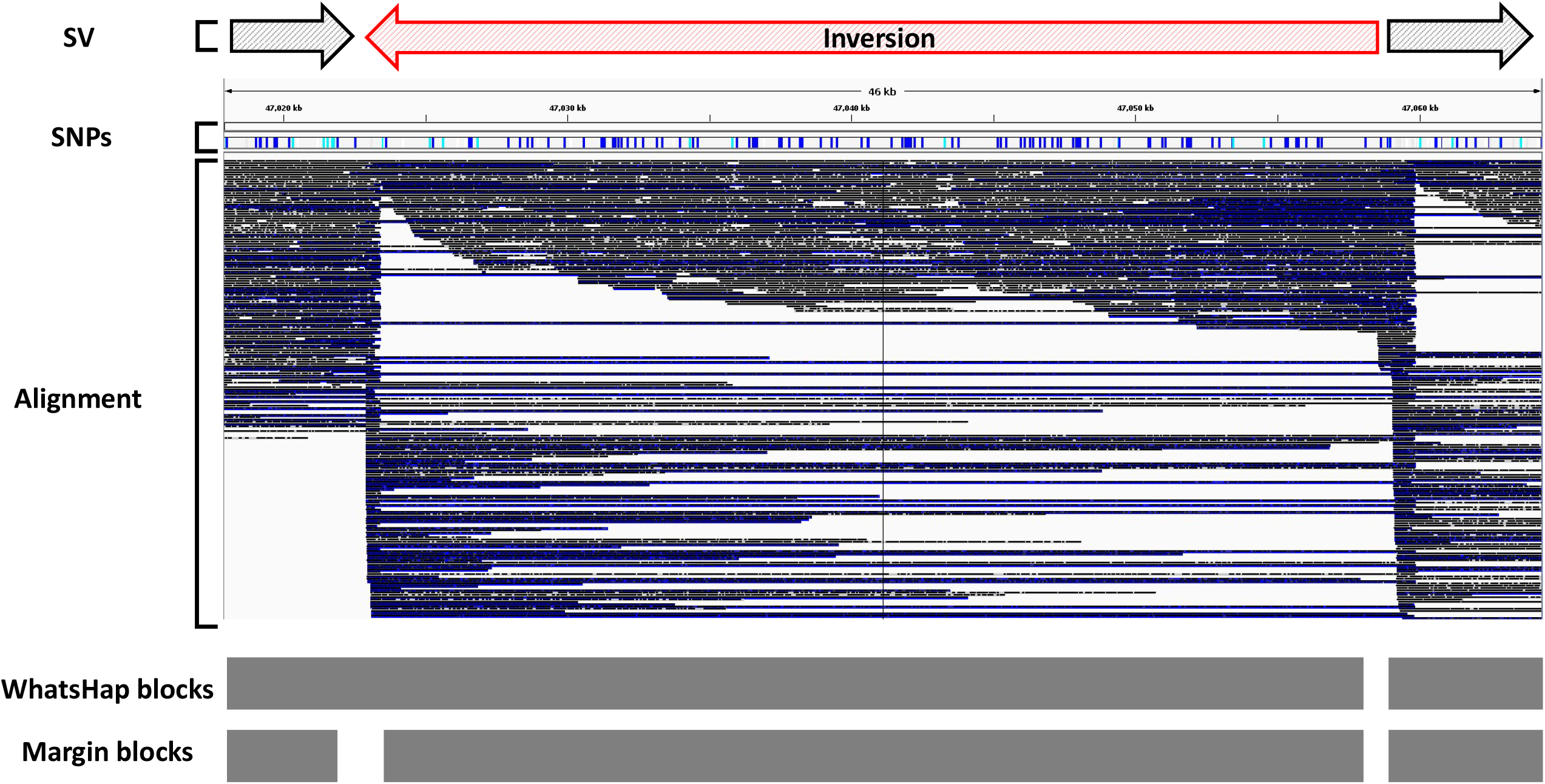
Illustration of broken read alignments and haplotype blocks around a ∼36kbp inversion. The long-read alignments are fragmented at the breakpoints of the inversion. Although the density of SNPs is high, the phased blocks of WhatsHap and Margin are both terminated at the inversion boundary.

Recently, several hybrid strategies have achieved chromosome-scale phasing by incorporating long-read sequencing with additional data. For instance, integrating long reads with proximity ligation (e.g., Hi-C) has reconstructed chromosome-scale haplotypes [12, 13]. In addition, the combination of long reads with single-cell strand-sequencing (Strand-seq) generated the first fully-phased human genome [14, 15]. Moreover, long-read sequencing of trios also demonstrated chromosome-scale phasing range via binning with Mendelian consistency [16, 17, 18]. However, the additional cost of these approaches is too high for population-scale phasing projects. Besides, the sequencing of trios is not always possible, and the trio-based method will lose novel variants in offspring [19, 20]. Consequently, the improvement of phasing range for long-read only sequencing is still in high demand for large-scale sequencing projects [21].

This paper presented an ultra-fast phasing algorithm, LongPhase, which simultaneously phases SNPs and SVs of a human genome in ∼10-20 minutes. We show that LongPhase can produce chromosome-scale haplotype blocks at compatible accuracy without the need for additional trios, chromosome-conformation, and single-cell sequencing.

## Results

### Co-phasing of SNPs and SVs by a two-stage phasing approach

LongPhase separately phases the human genome in two stages for maximizing efficiency (see method). We observed the majority of the human genome can be easily phased without using the entire reads. Only the remaining complex regions require full-lengthed reads. The utilization of long reads in two stages is much more efficient than previous approaches using the entire long reads from the beginning (e.g., WhatsHap and Margin). Initially, we formulate the SNPs and SVs as vertices in a directed acyclic graph (Figure 2). The edge stores the number of reads spanning the two variants as weight. The initial phasing computes the longest pairs of two disjoint paths in the graph, which is capable of tolerating a small amount of noises (e.g., sequencing errors and chimera).

**Figure 2.**
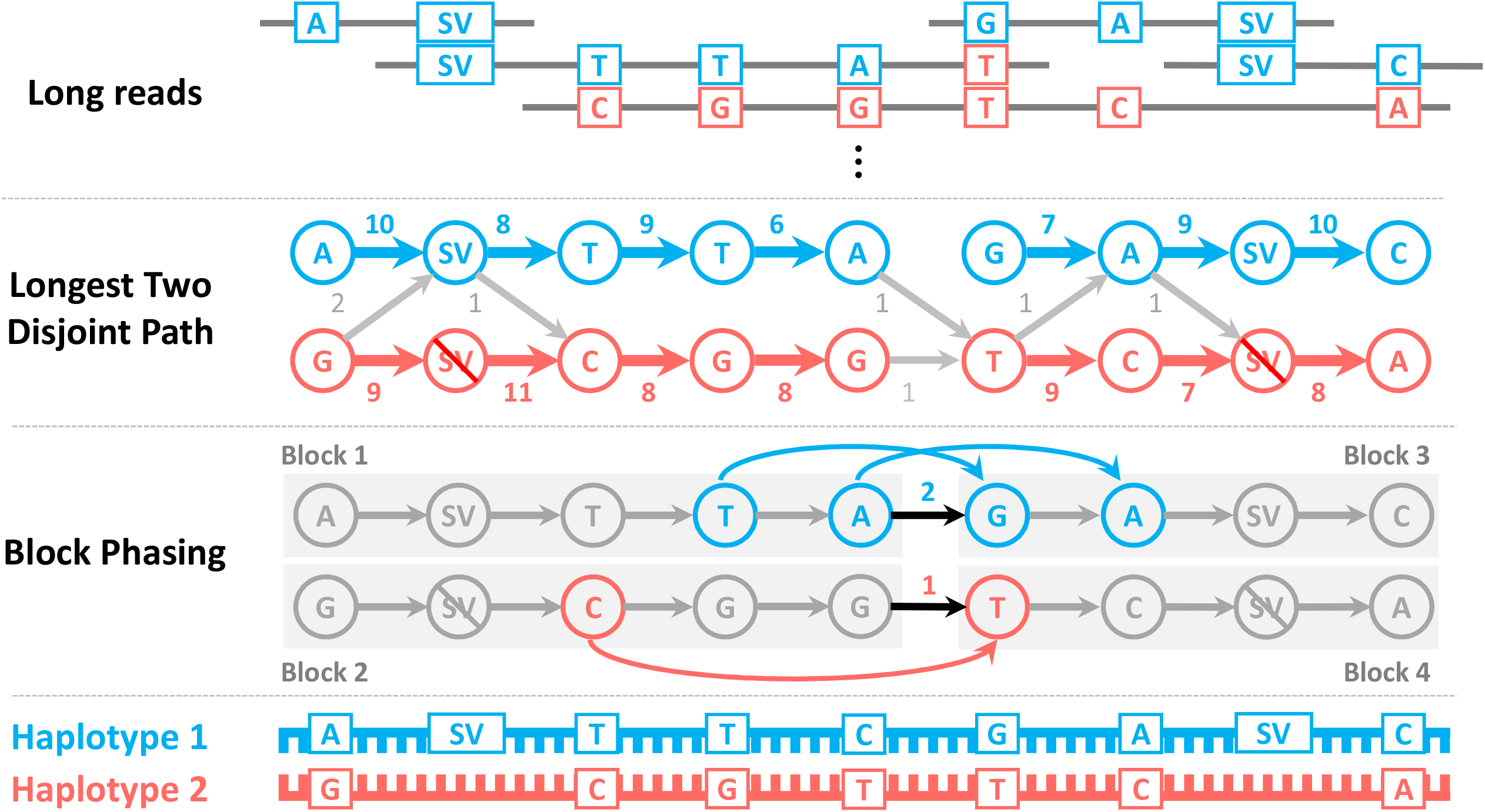
Illustration of the graph model in LongPhase for phasing SNPs and SVs. The SNPs and SVs are formulated as vertices in a graph, whereas the edges between two vertices stores the number of reads spanning two adjacent variants. The longest two-disjoint paths in the graph represent the two haplotype sequences supported by most reads.

The initial phasing breaks the two disjoint paths at loci without reads spanned or containing equally-weighted paths. These unphased loci are often due to false SNP/SV calls, spurious alignments, and/or systematic sequencing errors. However, they can be re-phased by the long-range linkage of distant variants in the long reads. Hence, in the second stage, the vertices become the initially-phased blocks, and the weights of the edges are the number of long reads spanning across SNPs/SVs in two adjacent blocks (e.g., two reads for joining Blocks 1 and 3 in Figure 2). Note that the size of the new graph is greatly reduced, because most SNPs/SVs have been phased and grouped into the same blocks. Finally, these broken blocks are phased again by finding the longest pairs of disjoint paths in the graph. By linking with the two disjoint paths found within each block, the two concatenated paths represent the paternal and maternal haplotypes supported by most reads.

The implemented program, LongPhase, can phase the entire human genome in ∼10-20 minutes on an eight-core (AMD) moderate machine (Figure 3(a)). Under the same environment and datasets, the state-of-the-art WhatsHap and Margin require 2-9 hours for phasing, which is 10-30 folds slower than LongPhase. Even though the running time of Margin can be reduced to 1-2 hours when moved to a 32-core machine, it is still six-fold slower than LongPhase. In particular, LongPhase can phase SNPs and SVs simultaneously, which is more efficient than previous twostage SV-phasing approaches [8, 7, 9, 10, 11].

**Figure 3.**
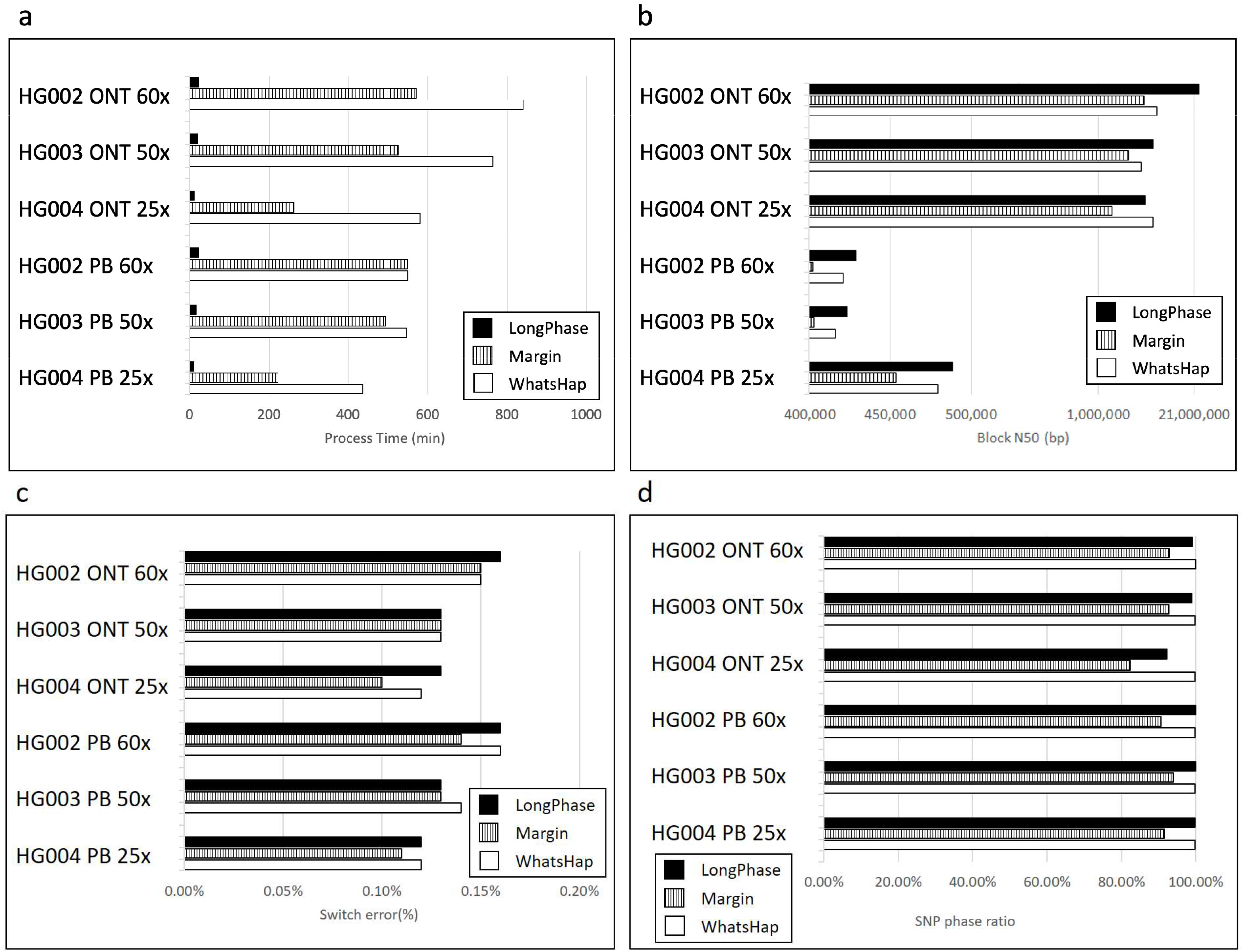
Comparison of LongPhase, Margin, and WhatsHap for (a) running time, (b) phasing contiguity, (c) switch errors, and (d) phasing percentage on HG002, HG003, and HG004 sequenced by Nanopore and PacBio at 25x, 50x, and 60x coverage.

### Comparison of accuracy and contiguity on Nanopore and PacBio sequencing

We compared LongPhase with WhatsHap and Margin on three public human genomes (HG002, HG003, and HG004) from the Genome in a Bottle (GIAB) consortium [22]. The three samples were sequenced by Nanopore and PacBio (HiFi) platforms. The sequencing reads are further down-sampled to 60x, 50x, and 25x coverage (Supplementary Table S1). The SNPs are called by the PEPPER-WhatsHap-DeepVariant pipelines [23], phased by the three programs, and evaluated by the GIAB benchmark (Supplementary Table S2).

In Nanopore sequencing, LongPhase can finish phasing in ∼10-20 minutes, while WhatsHap and Margin require 4-14 hours on the same datasets (Figure 3(a)). The N50 sizes of phased blocks by LongPhase (18-21Mbp) are significantly larger than those of WhatsHap (12-13Mbp) and Margin (3-10Mbp) (Figure 3(b), Supplementary Table S3). The largest haplotype block of LongPhase on HG002 reaches 93Mbp (∼51% of the chromosome 5), while those of WhatsHap and Margin are at most 47Mbp and 41Mbp, respectively. In terms of accuracy, Margin has relatively lower switch errors (0.1-0.15%) followed by WhatsHap (0.12-0.15%) and LongPhase (0.13-0.16%) (Figure 3(c)). However, most phasing errors of the three programs are found at the same loci. For instance, the 120 (out of 121) switch errors of Margin on HG002 are at the same loci as those of LongPhase and WhatsHap. Manual inspection by Integrative Genome Viewer (IGV) revealed that they are not errors (Supplementary Figure S1). Consequently, the phasing Benchmarks of GIAB may be still imperfect and the true phasing accuracy of all programs should be higher. Finally, the percentages of phased SNPs are the highest by WhatsHap (99.78-99.89%), which is followed by LongPhase (92.13-99.07%) and Margin (82.43-92.97%) (Figure 3(d)). Therefore, these results indicate that Margin trades phasing sensitivity for accuracy, while LongPhase and WhatsHap are more balanced at contiguity, sensitivity and accuracy.

In PacBio sequencing, LongPhase again runs faster (∼10-20 minutes) than the others (3-9 hours) (Figure 3(a)). The sizes of phased blocks of LongPhase are also larger (425kbp-487kbp) than those of WhatsHap (415-479kbp) and Margin (402-453kbp) (Figure 3(b)). All three programs exhibit similar switch errors: 0.12-0.16% for WhatsHap, 0.12-0.16% for LongPhase, and 0.11-0.14% for Margin (Figure 3(c)). Again, the SNP-phasing percentages of LongPhase (99.87-99.89%) and WhatsHap (99.81-99.86%) are significantly larger than those of Margin (90.60-93.94%) (Figure 3(d)). Although the phasing accuracy of PacBio reads is slightly higher than that of Nanopore, the phasing contiguity is much smaller than those of Nanopore. We note that the switch errors of Nanopore datasets are mainly due to systematic sequencing errors (see Discussion), which can be improved if using the latest basecaller and sequencing kit (e.g., Bonito and Q20EA). Hence, our results suggest that phasing by Nanopore ultra-long reads will provide larger contiguity with compatible accuracy.

### Chromosome-level phasing achieved by co-phasing SNPs and SVs

Next, we compare SNP/SV co-phasing with SNP-only phasing using LongPhase on the same datasets. 7,708-17,708 heterozygous SVs of HG002, HG003, and HG004, were called by Sniffles separately for Nanopore and PacBio sequencing (Supplementary tables S4) [24]. Deletions, insertions, duplications, inversions, and a few complex SVs are included in subsequent analysis (Supplementary Table S5). These SVs were co-phased with previously called SNPs by LongPhase. When tested on the 60x Nanopore dataset (HG002), co-phasing of SNPs and SVs reconstructs chromosome-scale haplotypes (N50=26Mbp), in which most of the chromosome arms are in a few blocks (Figure 4(a)) (Supplementary Table S6). For instance, the largest block (93Mbp) occupies ∼50% of chromosome 5. When compared with SNP-only phasing on HG002, HG003, and HG004 (Figure 4(b)), co-phasing of SNPs and SVs further improves the range of haplotype blocks at all sequencing depth, regardless of using PacBio or Nanopore. When focusing on SVs, *>*95% of SVs can be phased, and the phasing ratios mainly depend on the underlying sequencing depth (Figure 4(c)).

**Figure 4.**
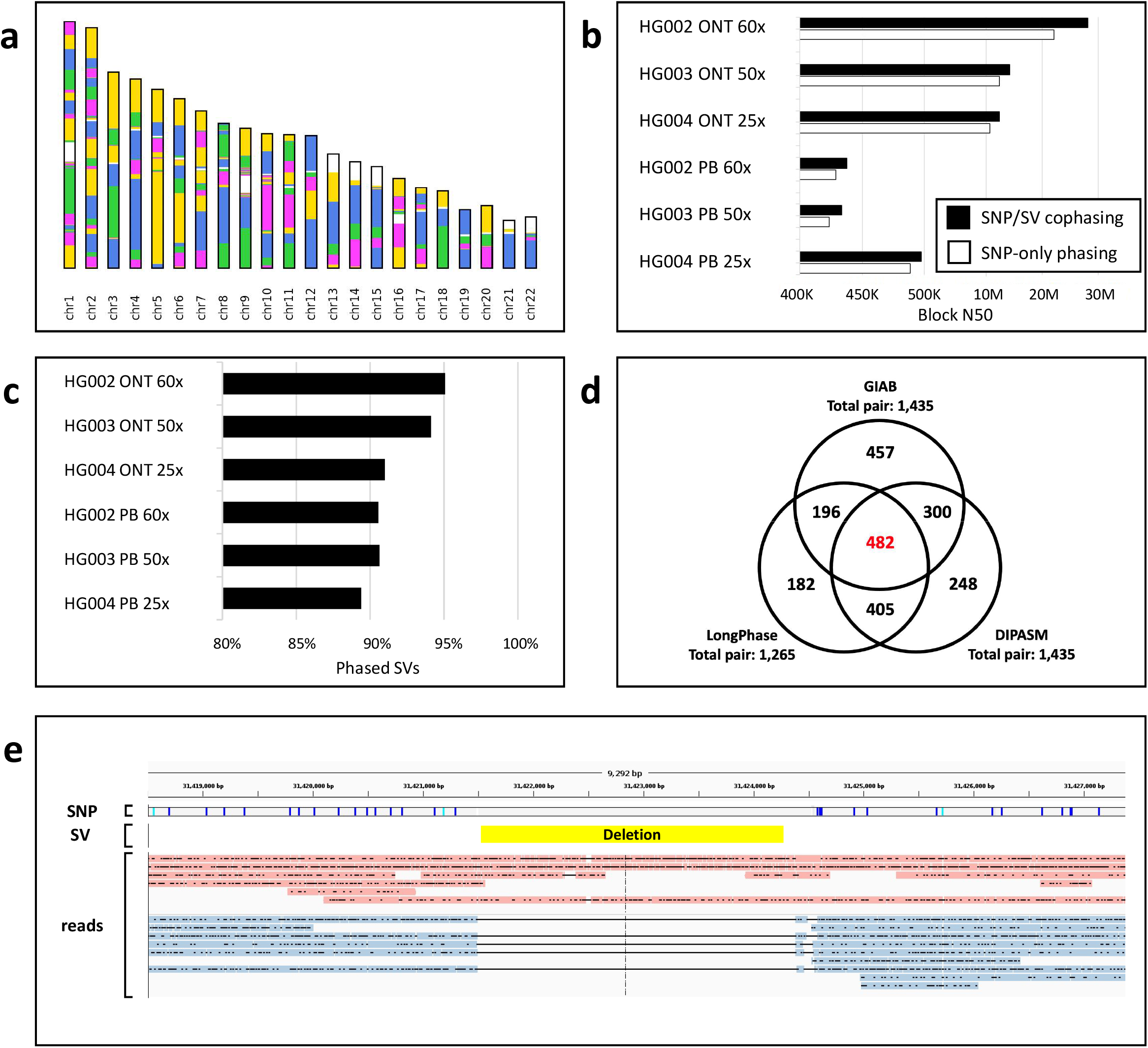
The contiguity and accuracy of SNP/SVs co-phasing by LongPhase. (a) Illustration of chromosome-scale phasing of LongPhasse on HG002. Each colored segment stands for a phased block, except for white blocks which represent centromere or acrocentric regions lacking of variants; (b) Comparison of N50 block sizes between SNP-only phasing and SNP/SV co-phasing on Nanopore and PacBio; (c) The ratios of phased SVs at different coverage sequenced by Nanopore and PacBio; (d) The Venn diagram of SV pairs uniquely or jointly phased by LongPhase, DipAsm, and GIAB; (e) Illustration of a ∼3kb deletion and SNPs cophased within the MHC region. The long reads are colored by the classified haplotypes, in which all the deletion reads are grouped in the same haplotype.

We assess the accuracy of phased SVs by comparing against two public datasets from GIAB and DipAsm [25, 26]. Specifically, the numbers of two adjacent SVs concordantly phased by any two methods are evaluated (see Methods). Although all methods agree on 482 pairs of phased SVs, each approach produces quite a few phased SVs inconsistent with others (e.g., 457 SV pairs only phased by GIAB). We observe LongPhase and DipAsm agree on most phasing pairs (i.e., 405) and have fewer numbers of uniquely-phased SVs (i.e., 182 and 248) when compared with GIAB. Furthermore, if using phased SVs of GIAB as ground truth, the switch error rates of LongPhase and DipAsm are ∼40%, but they reduce to ∼25% when compared against each other. Hence, these results suggest that the phased SVs in GIAB are less unreliable than SNPs. Because each method has a substantial amount of uniquely-phased SVs, a higher quality of benchmark is demanded for accuracy assessment. When switching the SV caller from Sniffles to CuteSV [27], the results are mainly the same (e.g., N50 = 26Mbp on HG002) (Supplementary Table S6), albeit the numbers of SVs called by Sniffles and CuteSV are quite different (e.g., 10,143 vs 15,176 deletions of Sniffles and CuteSV) (Supplementary Tables S5 and S7). This suggests that LongPhase is robust to the inconsistency of underlying SV detection algorithms (see Discussion).

Finally, we investigate the co-phasing of SNPs and SVs in the Major Histocompatibility Complex (MHC) region, which is highly polymorphic and essential for immune response. The entire MHC (∼4.9Mbp) of HG002 is phased by LongPhase and contained within a haplotype block of 30.8 Mbp (Figure 3(e)). In particular, one large hemizygous deletion (∼2.9kbp) is phased together with flanking 38,062 heterozygous SNPs in the block. By contrary, the MHC regions (of HG002) phased by WhatsHap and Margin are fragmented into six and four blocks, respectively. As a consequence, LongPhase is capable of phasing this complex region with frequent SNPs and SVs.

### Comparison of long-read only and hybrid phasing

Previously, chromosome-level phasing was mainly achieved by combining long-read sequencing with additional trios, proximity ligation (Hi-C), and/or single-cell Strand-seq. We compared LongPhase using only Nanopore reads with four hybrid methods on the same HG002 sample. Two used additional trios (Canu and Peregrine), one incorporated Hi-C (dipasm), and one utilized single-cell Strand-seq [26, 15, 18]. Table 1 lists the read length, sequencing depth, block N50, and switch errors of each method. LongPhase produces phasing contiguity (N50=26Mbp) similar to those using extra Hi-C (N50=25Mbp) and Strand-seq data (N50=26Mbp) (Table 1). On the other hand, the phasing contiguity of two trio-based methods is much smaller (e.g., 18Mbp by Canu and 16Mbp by Peregrine). We note that the long reads of these hybrid phasing approaches were produced by PacBio. As the read lengths of PacBio (N50=*<*13kb) are much shorter than those of Nanopore (N50=48kb), our results suggest that the increase in read length can compensate for additional trios, Hi-C, and strand-seq data.

**Table 1.**
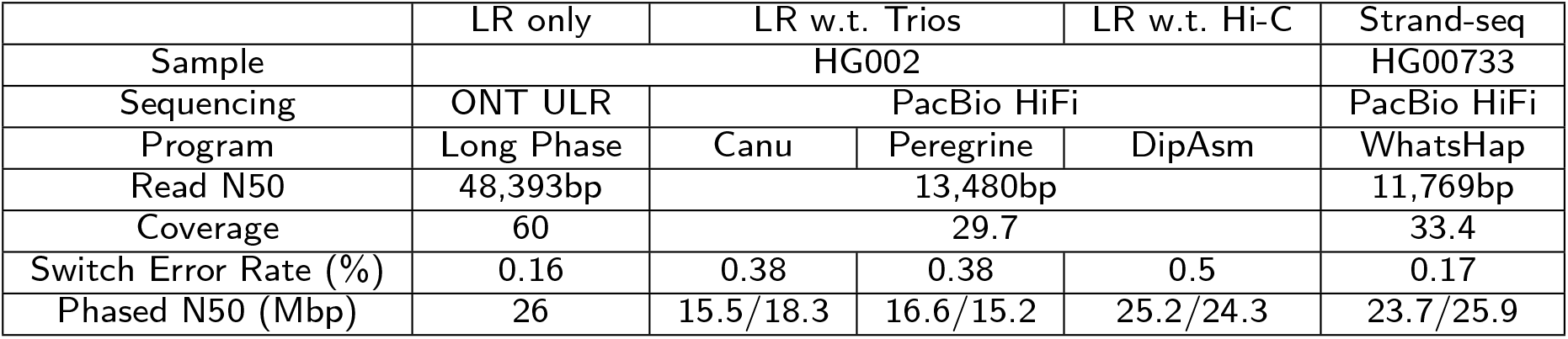
Comparison of LongPhase using only Nanopore ultra-long reads (ULR) with other hybrid phasing using long reads with additional trios, Hi-C, and Strand-seq data.

Furthermore, the error rate of LongPhase (0.16%) is on par with Strand-seq (0.17%), which is lower than the others utilizing trios and Hi-C sequencing (0.38-0.5%). The lower accuracy and contiguity of trio-based phasing might be owing to its limitations on heterozygous loci and/or novel variations generated in the offspring [26]. In summary, LongPhase with only Nanopore reads produces similar contiguity and accuracy compatible with the hybrid phasing using Strand-seq. In practice, the cost of single-cell Strand-seq is too expensive for population-scale phasing projects. Consequently, chromosome-scale phasing can be cost-effectively achieved solely by using Nanopore with LongPhase without the need for additional trios, proximity-ligation, or Strand-seq data.

## Discussion

This paper presents a new phasing program, LongPhase, which runs faster and generates much larger phasing blocks than WhatsHap and Margin at compatible accuracy. We observe LongPhase, WhatsHap, and Margin produced a similar set of switch errors at the same loci. Manual inspection by IGV reveals the errors are correct, suggesting the phasing benchmarks of GIAB are still imperfect. The remaining errors of LongPhase, manually inspected by IGV, are due to Nanopore systematic errors (e.g., methylated bases). These Nanopore systematic errors result in false SNP calls and/or wrong alleles, leading to switch errors during phasing. WhatsHap and Margin implemented different methods for correcting these systematic errors. For example, WhatsHap locally realigns reads at each locus for error correction. Margin re-estimates the probability of each allele of a variant using a hidden Markov model. By contrast, LongPhase still lacks a sophisticated error-correction module, though a few heuristics have been implemented (e.g., allele imputation, see Methods). In addition, we observe that Margin skips quite a few challenging loci for phasing. Consequently, it achieves higher accuracy yet sacrificing sensitivity and contiguity compared with LongPhase and WhatsHap. In the high-quality PacBio HiFi datasets, LongPhase exhibits the same accuracy as WhatsHap and Margin. Therefore, we expect the accuracy of LongPhase on Nanopore will be improved if using the high-quality kits and basecaller recently released (e.g., Q20+ kit and Bonito).

The accuracy of called SVs is often inferior in comparison with that of SNP calling (e.g., F1 *>* 0.999). In addition to false SV calls, we found that the many heterozygous SVs reported by Sniffles are homozygous and vice versa (Supplementary Figure S2) [24]. The newly developed SV caller, CuteSV, also infers inaccurate zygosity at many loci (Supplementary Figure S3) [27]. For example, in the Nanopore sequencing of HG002, Sniffles and CuteSV detected 42,650 and 51,237 SVs, respectively. However, only 15,532 SVs (30-36%) are concordantly called by the two programs. Among these high-confident SVs, the zygosity of 11,704 (75%) SVs, including 5,840 homozygous and 5,864 heterozygous variants, are consistently reported by each program. On the other hand, the zygosity of the remaining 3,828 (25%) SVs are inconsistent with each other, implying they are less reliable. These inaccurate SVs and zygosity slightly increase the switch error (by ∼0.01%) of LongPhase during co-phasing of SNPs and SVs. Therefore, the accuracy of SNP/SV co-phasing will rely on further improvement of underlying SV calling algorithms.

Although the accuracy of PacBio HiFi is higher than that of Nanopore, the contiguity of phased blocks is much smaller owing to a shorter read length (N50 = ∼15kbp). This is in contrast to the assembly of the Telomere-to-Telomere consortium [28], which constructs a highly contagious genome backbone starting with PacBio HiFi. We reason that genome assembly requires highly-accurate long reads for distinguishing highly-similar repeat polymorphisms. On the other hand, phasing demands reads with an overlapping length long enough to span at least two heterozygous variants. Because the distance between heterozygous variants is much larger, phasing benefits more from reads of longer length than those of higher quality. Recently, the ultra-long reads provided by Nanopore have further increased the length over 100kbp (e.g., N50=128kbp by Losse’s lab), significantly longer than the datasets used in this study (N50=48kbp). As a consequence, the phasing contiguity would be further improved with the increased read length. In addition, the accuracy gap between PacBio and Nanopore is closing with the recent release of Q20EA and Duplex reads (∼Q27). Consequently, LongPhase with Nanopore ultra-long reads is not only cost-effective compared with PacBio HiFi, but it can be also applied in sequencing projects at population scale without the need for additional trios, chromosome conformation, and Strand-seq data.

## Conclusion

We developed and implemented a novel long-read phasing algorithm, LongPhase, for simultaneously phasing small and large variants. The results indicated that LongPhase outperformed existing methods with much larger contiguity and faster running time at compatible accuracy. In particular, we showed that, in conjunction with Nanopore ultra-long reads, it can produce chromosome-level phasing without the need for additional trios, proximity ligation, and Strand-seq data. With the continuous improvement of read lengths and accuracy of Nanopore sequencing, we expect LongPhase may be a low-cost solution for population-scale phasing.

## Methods

### Data collection and variant calling

We downloaded public PacBio and Nanopore sequencing data of HG002, HG003, and HG004 from Genome in a Bottle (GIAB) [22]. The PacBio and Nanopore datasets were downsampled to 60x, 50x, and 25x (Supplementary Table S1). The long reads were aligned to the human genome (GRCh37) by minimap2 using specific parameters for Nanopore (-x map-ont) and PacBio HiFi (-x map-hifi) [29]. Subsequently, SNPs of each sample were called by the PEPPER-WhatsHap-DeepVariant pipeline [23], which produced over two millions of heterozygous SNPs (Supplementary Table S2). 7,787-17,708 and 11,677-25,813 heterozygous SVs were called by Sniffles (–num reads report) and CuteSV (–report readid –genotype), respectively (Supplementary Tables S4 and S5).

### Preprocessing of SNP phasing

When phasing SNPs, LongPhase takes the read alignment (in BAM format) and the SNPs (in VCF format) as input. The alignments with mapping quality scores less than 20 as well as all secondary alignments are ignored. All the heterozygous SNPs are sorted according to their coordinates on each chromosome. Subsequently, for each read overlapped with each heterozygous SNP, the allele carried by the read is extracted by parsing the corresponding CIGAR string in the alignment using htslib [30].

#### Removal of miscalled SNPs due to methylation

Two additional heuristics are implemented for reducing two types of Nanopore systematic errors: methylation and homopolymers. First, because the signals of electrical current of Nanopore are disturbed at methylated bases, these loci are often miscalled as SNPs by PEPPER-WhatsHap-DeepVariant and lead to phasing errors. We observe that, in true heterozygous SNPs, the major and minor alleles are usually found on reads from both strands. On the other hand, the miscalled SNPs due to methylation often exhibit strand bias. That is, it only carries the minor alleles on reads from either strand (Supplementary Figure S4). Hence, LongPhase requires that the two alleles of each heterozygous SNP must be presented on reads from both strands.

#### Imputation of missing alleles in homopolymers

Second, Nanopore sequencing fails to accurately estimate the length of a long homopolymer (e.g., long stretches of As). Because quite a few reads are wrongly base-called with insufficient length of homopolymers, the alleles of these SNPs are often missed (Supplementary Figure S5). Nevertheless, we observe these missing alleles (i.e., gaps in read alignments) are usually flanked with the correct one. Therefore, for SNPs within homopolymers of length at least three, the missing alleles are imputed by the flanking nucleotides.

### Preprocessing of SV Phasing

Sniffles and CuteSV can detect a variety of SVs, including deletions, insertions, duplications, inversions, translocations, and complex SVs. Manual inspection reveals that quite a few translocations outputted by these programs are inaccurate. Therefore, we skip translocations outputted by any program in downstream phasing. In addition, many SVs lead to fragmented alignments of long reads, in which many of them are tagged as supplementary alignments. These supplementary alignments with mapping quality at least 20 are included in subsequent phasing. The long reads supporting each SV are extracted by parsing the BAM alignment file. For SVs spanning an interval on the reference genome (e.g., deletions and inversions), the starting position of the SV is used as the representative locus in the subsequent graph formulation.

### A two-stage phasing approach

LongPhase develops a two-stage approach for efficiently phasing the entire human genome. We observe the majority of human genome can be easily phased without using the entire reads, while the remaining complex regions requires the usage of the whole long reads. This is much more efficient than previous approaches which utilize the long reads from the beginning (e.g., WhatsHap and Margin).

#### Initial phasing by solving the longest pairs of disjoint paths

Let *G* = (*V, E*) be a directed acyclic graph. The vertices *V* are the set of heterozygous SNPs or SVs topologically ordered according to their genomic positions. For each heterozygous SNP/SV, two vertices, denoted as *v* and *v′*, are created for representing the major and minor alleles. For any two adjacent SNP/SV variants, *v*_*i*_ and *v*_*i*+1_, four vertices are created and inter-connected by four edges, i.e., 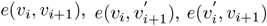 and 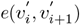 (Supplementary Figure S6). Each edge is associated with a weight which stores the number of reads supporting the connection of the two alleles.

Given *G*, we aim to find the longest pairs of two disjoint paths, where no common vertices and edges are shared in the two paths. Although the general version of the longest path problem is NP-hard, the directed acyclic graph can be solved in polynomial time by shortest-path algorithms applied on −*G* (i.e., change every weight to its negation). Theoretically, our problem can be solved by the Suurballe’s algorithm for finding edge disjoint shortest pairs in polynomial-time on −*G* [31]. But practically, the simplicity of the graph structure admits a simpler and faster greedy algorithm described below.

For any two adjacent SNPs/SVs, the longest pairs of disjoint paths of *G* can only be composed from one of the following two edge sets: *e*(*v*_*i*_, *v*_*i*+1_) with 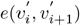 or 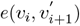 with 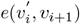. Hence, the larger weight of the two sets will contribute to the optimal solution. Because the optimal solution of *G* is composed of optimal solutions of subgraphs between any two adjacent SNPs/SVs, a greedy algorithm can be applied. Specifically, by following the topological order, the longest pairs of disjoint paths are iteratively constructed by picking the edge set of the larger sum for any two adjacent SNPs/SVs (Supplementary Figure S7). The union of edge sets with larger weight between each two adjacent SNPs/SVs is thus the longest pairs of disjoint paths. Note that we break the two disjoint paths at loci without reads spanned or containing equally-weighted paths. Since *G* is a sparse graph, the running time of this algorithm is *O*(*V*).

#### Block phasing by long-range linkage

The initially-phased blocks are often broken by false SNP/SV calls, repeat artifacts, and/or elevated sequencing errors, which create either no or false edges between two adjacent vertices in the initial graph. Because the initial phasing only extracts short linkage between adjacent variants from reads, the algorithm is very efficient but vulnerable to these noises. Fortunately, most of these noises can be removed by using the long-range linkage between distant variants in the long reads. Specifically, each haplotype of the initially-phased blocks are formulated as vertices, and the edges represent there are reads spanning across variants of two haplotypes (Supplementary Figure S8). Similarly, the edge weight is the number of reads supporting the haplotype connection. For improving efficiency, at most ten SNPs/SVs closest to the boundary of each block are used for edge creation and weight calculation. Note that the size of the new graph is greatly reduced, as most SNPs/SVs have been phased and collapsed into the same blocks. Finally, the longest pairs of two disjoint paths are computed by the the same greedy algorithm. By combining with the previously computed disjoint paths within each block, the two concatenated disjoint paths represent the paternal and maternal haplotypes supported by most amount of reads.

To implement the execution time of the software, we don’t need to check all the reads shared by any two initially-phased blocks. Since an average ultra-long read will only contain 19 heterozygous SNPs, we only need to check the SNPs before and after the two blocks. The number of SNPs we checked was higher than average. We check 15 SNPs in each block, totaling 30 SNPs. Since we use the primary and supplementary alignment of a read at the same time, we may encounter chimeric reads. This will cause two blocks that should not be connected to be phased together. By default, two blocks with a distance of more than 300kbp will not be phasing.

### Evaluations of SNP and SV phasing accuracy

To validate the accuracy of phased SNPs of each program, the phased haplotypes are compared with GIAB phased benchmark, HG002 NISTv4.2.1, HG003 NISTv3.3, and HG004 NISTv3.3 using “WhatsHap compare” [2]. The phased SNPs of the three benchmarks are 275,057, 234,719, and 240,484, respectively. The switch errors were computed by WhatsHap compare.

Two public phased SVs of HG002 are used for validating the accuracy of SV phasing. The first one is downloaded from GIAB’s Tier v0.6 (PBHP GT) [25]. The second one is retrieved from the haplotype-resolved assembly by DipAsm (NA24385-denovo.dip.vcf.gz) [26], in which the haplotype-resolved SVs were called by DipCall [32]. We found that the phased SVs in GIAB are also in silico prediction and largely inconsistent with those of LongPhase and DipAsm/DipCall. Therefore, we computed the number of SV pairs concordantly or uniquely phased by each program. However, the comparison of SVs of these programs are further complicated by non-overlapping SVs generated by different callers. We use TRUVARI for identifying 1,457 SVs commonly overlapped in GIAB, DipAsm, and LongPhase [33]. By focusing on these commonly-overlapped SVs, we can then compute the numbers of common or uniquely pairs of phasing SVs outputted by each program.

## Acknowledgement

The authors would like to thank Dr. Chia-Lin Wei of the Jackson Laboratory for not only motivating but also helping us along the way. This work won’t be possible without her kind support.

## Author’s contributions

YTH conceived the study. JHL and YTH designed the algorithms and experiments. JHL is in charge of most implementation. LCC and SQY helped implement the parsing module and multi-threading parallelism. YTH and JHL wrote the manuscript. All authors agreed on the the final version of the manuscript.

## Ethical approval and consent to participate

Not applicable.

## Consent to publication

Not applicable.

## Availability of Data and Materials

LongPhase is freely available at https://github.com/twolinin/LongPhase/.

## Additional information

### Additional file 1

Additional file 1 includes Supplementary Figs. S1-8. Fig. S1. Comparison of phasing results of LongPhase, WhatsHap, Margin, and GIAB. Fig. S2. An example of homozygous SVs miscalled as heterozygous genotypes by Sniffles. Fig. S3. An example of homozygous SVs miscalled as heterozygous genotypes by CuteSV. Fig. S4. Strand bias of miscalled SNPs due to methylation. Fig. S5. Illustration of Nanopore systematic errors in homopolymer regions. Fig. S6. Illustration of the graph model used by LongPhase. Fig. S7. Illustration of the greedy algorithm for finding the longest disjoint paths. Fig. S8. Illustration of block phasing by long-range linkage.

### Additional file 2

Additional file 2 includes Supplementary Tables S1-7. Table S1. Sequencing statistics of Nanopore and PacBio on HG002, HG003, and HG004. Table S2. PEPPER SNP calling statistics of HG002, HG003, and HG004. Table S3. SNP-only phasing statistics of LongPhase, Margin, and WhatsHap. Table S4. Statistics of SVs called by Sniffles and CuteSV on HG002, HG003, and HG004. Table S5. Types of heterozygous SVs called by Sniffles. Table S6. Statistics of SNP/SV co-phasing by LongPhase. Table S7. Types of heterozygous SVs called by CuteSV.

## References

1. Garg, S.: Computational methods for chromosome-scale haplotype reconstruction. Genome Biol. 22(1), 101 (2021)

2. Patterson, M., Marschall, T., Pisanti, N., van Iersel, L., Stougie, L., Klau, G.W., Schönhuth, A.: WhatsHap: Weighted haplotype assembly for Future-Generation sequencing reads. J. Comput. Biol. 22(6), 498–509 (2015)

3. Martin, M., Patterson, M., Garg, S., Fischer, S., Pisanti, N., Klau, G.W., Schöenhuth, A., Marschall, T.: WhatsHap: fast and accurate read-based phasing (2016)

4. Edge, P., Bafna, V., Bansal, V.: HapCUT2: robust and accurate haplotype assembly for diverse sequencing technologies. Genome Res. 27(5), 801–812 (2017)

5. Ebler, J., Haukness, M., Pesout, T., Marschall, T., Paten, B.: Haplotype-aware diplotyping from noisy long reads. Genome Biol. 20(1), 116 (2019)

6. Shafin, K., Pesout, T., Chang, P.-C., Nattestad, M., Kolesnikov, A., Goel, S., Baid, G., Eizenga, J.M., Miga, K.H., Carnevali, P., Jain, M., Carroll, A., Paten, B.: Haplotype-aware variant calling enables high accuracy in nanopore long-reads using deep neural networks (2021)

7. Cretu Stancu, M., van Roosmalen, M.J., Renkens, I., Nieboer, M.M., Middelkamp, S., de Ligt, J., Pregno, G., Giachino, D., Mandrile, G., Espejo Valle-Inclan, J., Korzelius, J., de Bruijn, E., Cuppen, E., Talkowski, M.E., Marschall, T., de Ridder, J., Kloosterman, W.P.: Mapping and phasing of structural variation in patient genomes using nanopore sequencing. Nat. Commun. 8(1), 1326 (2017)

8. Garg, S., Fungtammasan, A., Carroll, A., Chou, M., Schmitt, A., Zhou, X., Mac, S., Peluso, P., Hatas, E., Ghurye, J., Maguire, J., Mahmoud, M., Cheng, H., Heller, D., Zook, J.M., Moemke, T., Marschall, T., Sedlazeck, F.J., Aach, J., Chin, C.-S., Church, G.M., Li, H.: Chromosome-scale, haplotype-resolved assembly of human genomes. Nat. Biotechnol. 39(3), 309–312 (2021)

9. Rodriguez, O.L., Ritz, A., Sharp, A.J., Bashir, A.: MsPAC: a tool for haplotype-phased structural variant detection. Bioinformatics 36(3), 922–924 (2020)

10. Chaisson, M.J.P., Sanders, A.D., Zhao, X., Malhotra, A., Porubsky, D., Rausch, T., Gardner, E.J., Rodriguez, O.L., Guo, L., Collins, R.L., Fan, X., Wen, J., Handsaker, R.E., Fairley, S., Kronenberg, Z.N., Kong, X., Hormozdiari, F., Lee, D., Wenger, A.M., Hastie, A.R., Antaki, D., Anantharaman, T., Audano, P.A., Brand, H., Cantsilieris, S., Cao, H., Cerveira, E., Chen, C., Chen, X., Chin, C.-S., Chong, Z., Chuang, N.T., Lambert, C.C., Church, D.M., Clarke, L., Farrell, A., Flores, J., Galeev, T., Gorkin, D.U., Gujral, M., Guryev, V., Heaton, W.H., Korlach, J., Kumar, S., Kwon, J.Y., Lam, E.T., Lee, J.E., Lee, J., Lee, W.-P., Lee, S.P., Li, S., Marks, P., Viaud-Martinez, K., Meiers, S., Munson, K.M., Navarro, F.C.P., Nelson, B.J., Nodzak, C., Noor, A., Kyriazopoulou-Panagiotopoulou, S., Pang, A.W.C., Qiu, Y., Rosanio, G., Ryan, M., Stütz, A., Spierings, D.C.J., Ward, A., Welch, A.E., Xiao, M., Xu, W., Zhang, C., Zhu, Q., Zheng-Bradley, X., Lowy, E., Yakneen, S., McCarroll, S., Jun, G., Ding, L., Koh, C.L., Ren, B., Flicek, P., Chen, K., Gerstein, M.B., Kwok, P.-Y., Lansdorp, P.M., Marth, G.T., Sebat, J., Shi, X., Bashir, A., Ye, K., Devine, S.E., Talkowski, M.E., Mills, R.E., Marschall, T., Korbel, J.O., Eichler, E.E., Lee, C.: Multi-platform discovery of haplotype-resolved structural variation in human genomes. Nat. Commun. 10(1), 1784 (2019)

11. Li, H., Bloom, J.M., Farjoun, Y., Fleharty, M., Gauthier, L., Neale, B., MacArthur, D.: A synthetic-diploid benchmark for accurate variant-calling evaluation. Nat. Methods 15(8), 595–597 (2018)

12. Kronenberg, Z.N., Rhie, A., Koren, S., Concepcion, G.T., Peluso, P., Munson, K.M., Porubsky, D., Kuhn, K., Mueller, K.A., Low, W.Y., Hiendleder, S., Fedrigo, O., Liachko, I., Hall, R.J., Phillippy, A.M., Eichler, E.E., Williams, J.L., Smith, T.P.L., Jarvis, E.D., Sullivan, S.T., Kingan, S.B.: Extended haplotype-phasing of long-read de novo genome assemblies using Hi-C. Nat. Commun. 12(1), 1935 (2021)

13. Tourdot, R.W., Brunette, G.J., Pinto, R.A., Zhang, C.-Z.: Determination of complete chromosomal haplotypes by bulk DNA sequencing. Genome Biol. 22(1), 139 (2021)

14. Ebert, P., Audano, P.A., Zhu, Q., Rodriguez-Martin, B., Porubsky, D., Bonder, M.J., Sulovari, A., Ebler, J., Zhou, W., Serra Mari, R., Yilmaz, F., Zhao, X., Hsieh, P., Lee, J., Kumar, S., Lin, J., Rausch, T., Chen, Y., Ren, J., Santamarina, M., Höps, W., Ashraf, H., Chuang, N.T., Yang, X., Munson, K.M., Lewis, A.P., Fairley, S., Tallon, L.J., Clarke, W.E., Basile, A.O., Byrska-Bishop, M., Corvelo, A., Evani, U.S., Lu, T.-Y., Chaisson, M.J.P., Chen, J., Li, C., Brand, H., Wenger, A.M., Ghareghani, M., Harvey, W.T., Raeder, B., Hasenfeld, P., Regier, A.A., Abel, H.J., Hall, I.M., Flicek, P., Stegle, O., Gerstein, M.B., Tubio, J.M.C., Mu, Z., Li, Y.I., Shi, X., Hastie, A.R., Ye, K., Chong, Z., Sanders, A.D., Zody, M.C., Talkowski, M.E., Mills, R.E., Devine, S.E., Lee, C., Korbel, J.O., Marschall, T., Eichler, E.E.: Haplotype-resolved diverse human genomes and integrated analysis of structural variation. Science 372(6537) (2021)

15. Porubsky, D., Ebert, P., Audano, P.A., Vollger, M.R., Harvey, W.T., Marijon, P., Ebler, J., Munson, K.M., Sorensen, M., Sulovari, A., Haukness, M., Ghareghani, M., Human Genome Structural Variation Consortium, Lansdorp, P.M., Paten, B., Devine, S.E., Sanders, A.D., Lee, C., Chaisson, M.J.P., Korbel, J.O., Eichler, E.E., Marschall, T.: Fully phased human genome assembly without parental data using single-cell strand sequencing and long reads. Nat. Biotechnol. 39(3), 302–308 (2021)

16. Cheng, H., Concepcion, G.T., Feng, X., Zhang, H., Li, H.: Haplotype-resolved de novo assembly using phased assembly graphs with hifiasm. Nat. Methods 18(2), 170–175 (2021)

17. Koren, S., Rhie, A., Walenz, B.P., Dilthey, A.T., Bickhart, D.M., Kingan, S.B., Hiendleder, S., Williams, J.L., Smith, T.P.L., Phillippy, A.M.: Complete assembly of parental haplotypes with trio binning. BioRxiv (2018)

18. Wenger, A.M., Peluso, P., Rowell, W.J., Chang, P.-C., Hall, R.J., Concepcion, G.T., Ebler, J., Fungtammasan, A., Kolesnikov, A., Olson, N.D., Töpfer, A., Alonge, M., Mahmoud, M., Qian, Y., Chin, C.-S., Phillippy, A.M., Schatz, M.C., Myers, G., DePristo, M.A., Ruan, J., Marschall, T., Sedlazeck, F.J., Zook, J.M., Li, H., Koren, S., Carroll, A., Rank, D.R., Hunkapiller, M.W.: Accurate circular consensus long-read sequencing improves variant detection and assembly of a human genome. Nat. Biotechnol. 37(10), 1155–1162 (2019)

19. Koren, S., Rhie, A., Walenz, B.P., Dilthey, A.T., Bickhart, D.M., Kingan, S.B., Hiendleder, S., Williams, J.L., Smith, T.P.L., Phillippy, A.M.: De novo assembly of haplotype-resolved genomes with trio binning. Nat. Biotechnol. (2018)

20. Garg, S., Aach, J., Li, H., Sebenius, I., Durbin, R., Church, G.: A haplotype-aware de novo assembly of related individuals using pedigree sequence graph. Bioinformatics 36(8), 2385–2392 (2020)

21. De Coster, W., Weissensteiner, M.H., Sedlazeck, F.J.: Towards population-scale long-read sequencing. Nat. Rev. Genet. (2021)

22. Zook, J.M., Catoe, D., McDaniel, J., Vang, L., Spies, N., Sidow, A., Weng, Z., Liu, Y., Mason, C.E., Alexander, N., Henaff, E., McIntyre, A.B.R., Chandramohan, D., Chen, F., Jaeger, E., Moshrefi, A., Pham, K., Stedman, W., Liang, T., Saghbini, M., Dzakula, Z., Hastie, A., Cao, H., Deikus, G., Schadt, E., Sebra, R., Bashir, A., Truty, R.M., Chang, C.C., Gulbahce, N., Zhao, K., Ghosh, S., Hyland, F., Fu, Y., Chaisson, M., Xiao, C., Trow, J., Sherry, S.T., Zaranek, A.W., Ball, M., Bobe, J., Estep, P., Church, G.M., Marks, P., Kyriazopoulou-Panagiotopoulou, S., Zheng, G.X.Y., Schnall-Levin, M., Ordonez, H.S., Mudivarti, P.A., Giorda, K., Sheng, Y., Rypdal, K.B., Salit, M.: Extensive sequencing of seven human genomes to characterize benchmark reference materials. Sci Data 3, 160025 (2016)

23. Cook, D.: Improving Variant Calling using Haplotype Information. https://google.github.io/deepvariant/posts/2021-02-08-the-haplotype-channel/. Accessed: 2021-5-21 (2021)

24. Sedlazeck, F.J., Rescheneder, P., Smolka, M., Fang, H., Nattestad, M., von Haeseler, A., Schatz, M.C.: Accurate detection of complex structural variations using single-molecule sequencing. Nat. Methods 15(6), 461–468 (2018)

25. Zook, J.M., Hansen, N.F., Olson, N.D., Chapman, L., Mullikin, J.C., Xiao, C., Sherry, S., Koren, S., Phillippy, A.M., Boutros, P.C., et al.: A robust benchmark for detection of germline large deletions and insertions. Nature biotechnology 38(11), 1347–1355 (2020)

26. Garg, S., Fungtammasan, A., Carroll, A., Chou, M., Schmitt, A., Zhou, X., Mac, S., Peluso, P., Hatas, E., Ghurye, J., et al.: Chromosome-scale, haplotype-resolved assembly of human genomes. Nature biotechnology 39(3), 309–312 (2021)

27. Jiang, T., Liu, Y., Jiang, Y., Li, J., Gao, Y., Cui, Z., Liu, Y., Liu, B., Wang, Y.: Long-read-based human genomic structural variation detection with cuteSV. Genome Biol. 21(1), 189 (2020)

28. Nurk, S., Koren, S., Rhie, A., Rautiainen, M., Bzikadze, A.V., Mikheenko, A., Vollger, M.R., Altemose, N., Uralsky, L., Gershman, A., Aganezov, S., Hoyt, S.J., Diekhans, M., Logsdon, G.A., Alonge, M., Antonarakis, S.E., Borchers, M., Bouffard, G.G., Brooks, S.Y., Caldas, G.V., Cheng, H., Chin, C.-S., Chow, W., de Lima, L.G., Dishuck, P.C., Durbin, R., Dvorkina, T., Fiddes, I.T., Formenti, G., Fulton, R.S., Fungtammasan, A., Garrison, E., Grady, P.G.S., Graves-Lindsay, T.A., Hall, I.M., Hansen, N.F., Hartley, G.A., Haukness, M., Howe, K., Hunkapiller, M.W., Jain, C., Jain, M., Jarvis, E.D., Kerpedjiev, P., Kirsche, M., Kolmogorov, M., Korlach, J., Kremitzki, M., Li, H., Maduro, V.V., Marschall, T., McCartney, A.M., McDaniel, J., Miller, D.E., Mullikin, J.C., Myers, E.W., Olson, N.D., Paten, B., Peluso, P., Pevzner, P.A., Porubsky, D., Potapova, T., Rogaev, E.I., Rosenfeld, J.A., Salzberg, S.L., Schneider, V.A., Sedlazeck, F.J., Shafin, K., Shew, C.J., Shumate, A., Sims, Y., Smit, A.F.A., Soto, D.C., Sović, I., Storer, J.M., Streets, A., Sullivan, B.A., Thibaud-Nissen, F., Torrance, J., Wagner, J., Walenz, B.P., Wenger, A., Wood, J.M.D., Xiao, C., Yan, S.M., Young, A.C., Zarate, S., Surti, U., McCoy, R.C., Dennis, M.Y., Alexandrov, I.A., Gerton, J.L., O’Neill, R.J., Timp, W., Zook, J.M., Schatz, M.C., Eichler, E.E., Miga, K.H., Phillippy, A.M.: The complete sequence of a human genome (2021)

29. Li, H.: Minimap2: pairwise alignment for nucleotide sequences. Bioinformatics 34(18), 3094–3100 (2018)

30. Bonfield, J.K., Marshall, J., Danecek, P., Li, H., Ohan, V., Whitwham, A., Keane, T., Davies, R.M.: HTSlib: C library for reading/writing high-throughput sequencing data. GigaScience 10(2) (2021). doi:10.1093/gigascience/giab007.giab007. https://academic.oup.com/gigascience/article-pdf/10/2/giab007/36332285/giab007.pdf

31. Suurballe, J.W.: Disjoint paths in a network. Networks 4(2), 125–145 (1974). doi:10.1002/net.3230040204. https://onlinelibrary.wiley.com/doi/pdf/10.1002/net.3230040204

32. Li, H., Bloom, J.M., Farjoun, Y., Fleharty, M., Gauthier, L., Neale, B., MacArthur, D.: A synthetic-diploid benchmark for accurate variant-calling evaluation. Nature methods 15(8), 595–597 (2018)

33. Zook, J.M., Hansen, N.F., Olson, N.D., Chapman, L.M., Mullikin, J.C., Xiao, C., Sherry, S., Koren, S., Phillippy, A.M., Boutros, P.C., et al.: A robust benchmark for germline structural variant detection. BioRxiv, 664623 (2019)

