## Additional File 1 for "LongPhase: an ultra-fast chromosome-scale phasing algorithm for small and large variants"

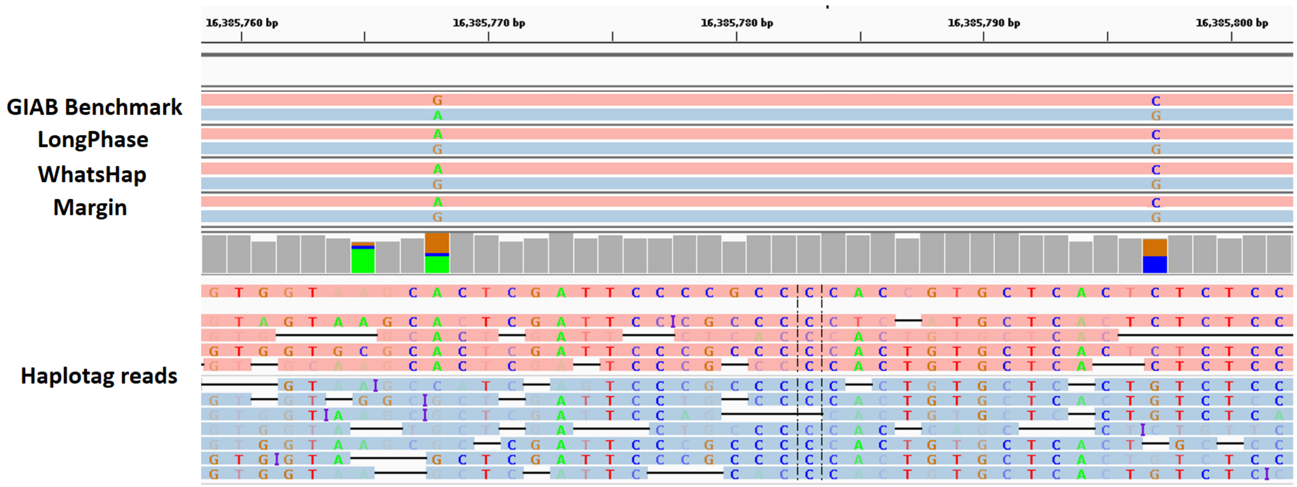


**Fig. S1:** The phased results of LongPhase, WhatsHap, and Margin are consistent with each other (i.e., AC/GG), while those of the GIAB benchmark disagrees with all the others (i.e., GC/AG). The raw reads in the bottom of IGV agrees with the results of three programs and disagree with those of GIAB.

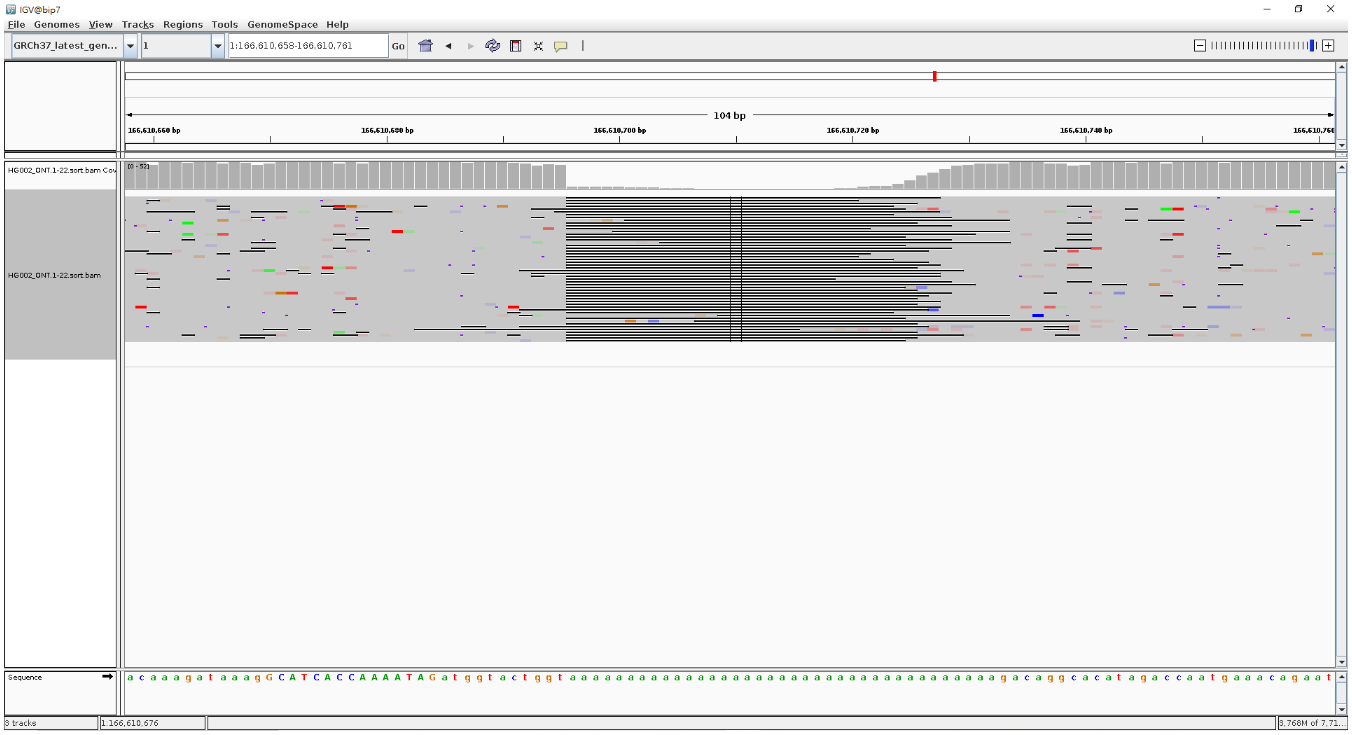

**Fig. S2:** An example of homozygous SVs miscalled as heterozygous genotypes by Sniffles.

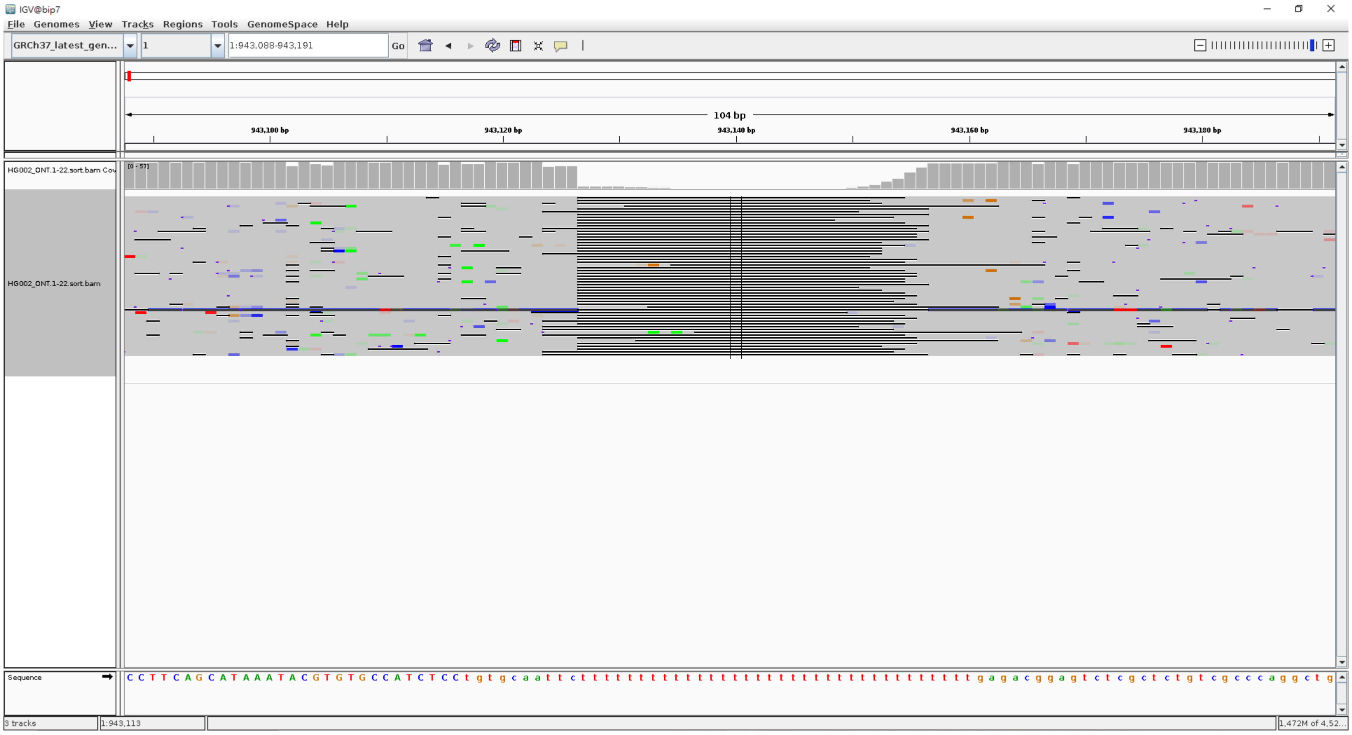


**Fig. S3:** An example of homozygous SVs miscalled as heterozygous genotypes by CuteSV.

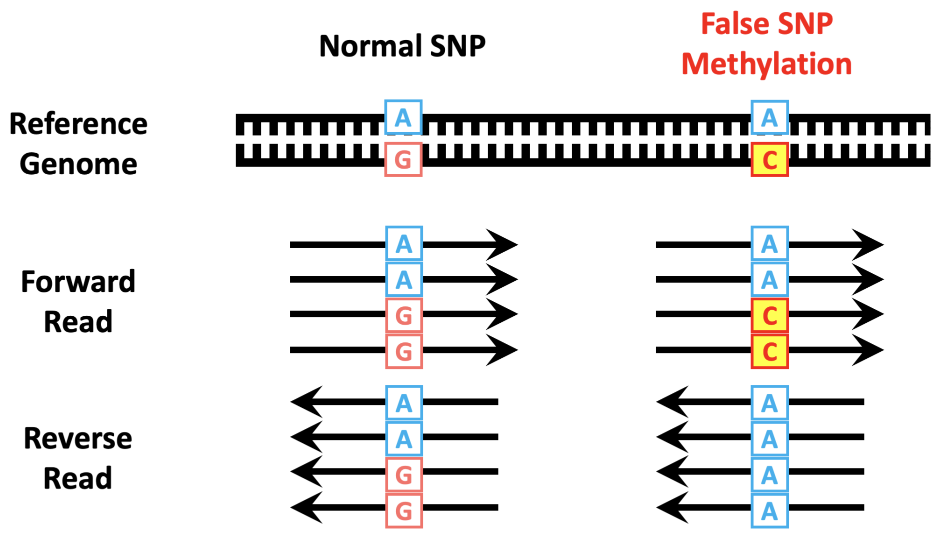


**Fig. S4:** Strand bias of methylation-induced SNPs. Ordinary heterozygous SNPs possess the major and minor alleles on both the forward and reverse strands. On the other hand, the miscalled SNPs due to methylation often carry the minor alleles on only one strand.

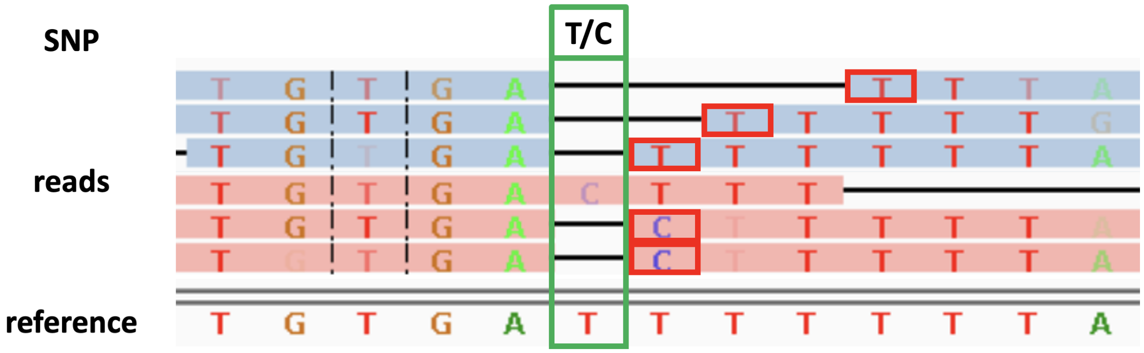


**Fig. S5:** Illustration of Nanopore systematic errors in homopolymer regions. The correct alleles, C and T, are right-shifted in five read alignments owing to shorter homopolymers incorrectly basecalled. LongPhase imputes the missed alleles in homopolymers using the right-shifted base.

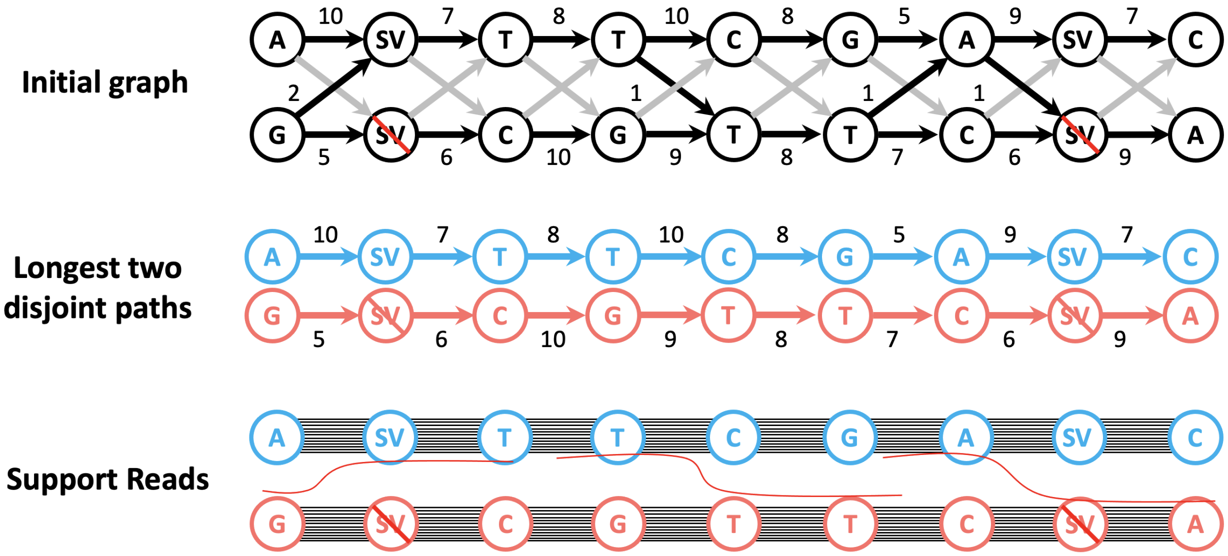


**Fig. S6:** Illustration of the graph model used by LongPhase. The black edges store the number of long reads supporting the linkage of the two alleles. The adjacent vertices without supporting reads are added with grey edges of zero weights. The longest pairs of disjoint paths are computed by a greedy algorithm, which represents the two haplotypes supported by most reads.

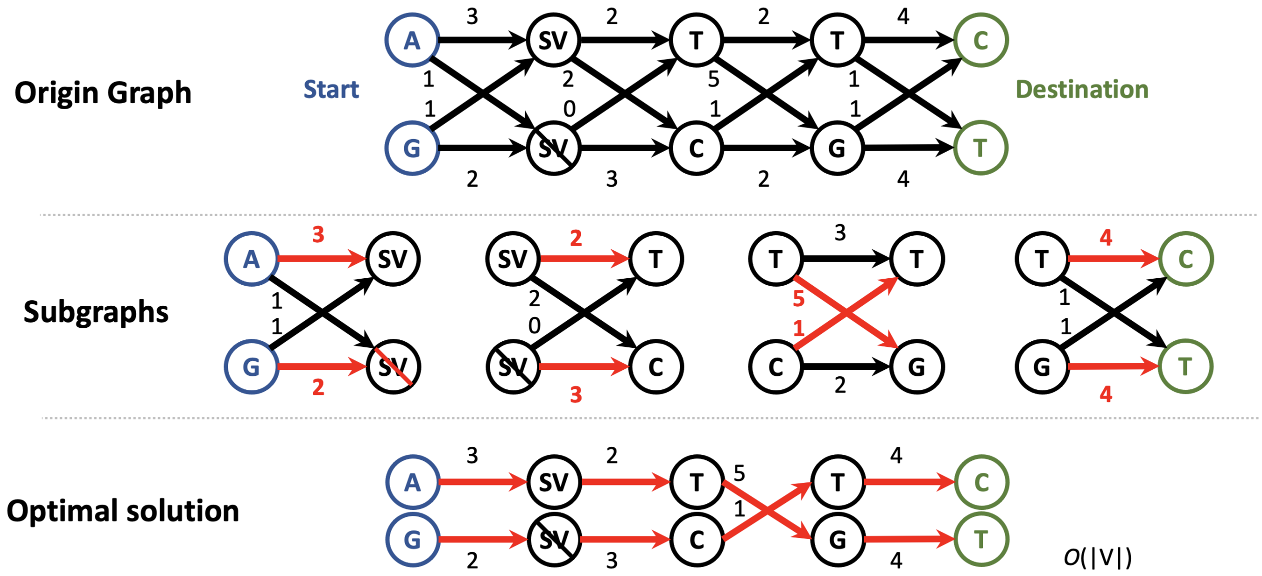


**Fig. S7:** Illustration of the greedy algorithm for finding the longest disjoint paths. For any two adjacent variant loci, because there are only two possible sets of interconnecting edges, the algorithm greedily selects the set with larger weight according to the topological order of vertices.

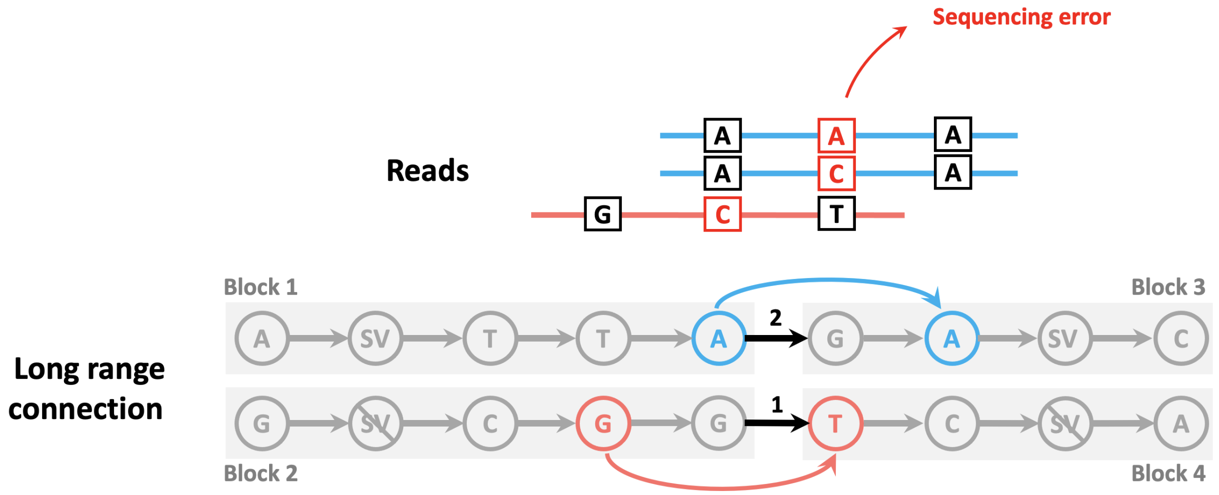


**Fig. S8:** Illustration of block phasing by long-range linkage. In the initial phasing, a few erroneous edges lead to termination of phasing. In the block-phasing stage, the long-range linkage between distant variants of two adjacent blocks are extracted from long reads, which will be used to joint the broken blocks into a larger one.
