## Additional File 2 for "LongPhase: an ultra-fast chromosome-scale phasing algorithm for small and large variants"

**Table S1:** Sequencing statistics of Nanopore and PacBio on HG002, HG003, and HG004.

| Sequence | Sample | Error rate | No. of Read | N50 | Total Base | Coverage |
|---|---|---|---|---|---|---|
| Nanopore ultra-long | HG002 | 11% | 19,328,993 | 48,393 | 156,500,000,000 | 58 |
| Nanopore ultra-long | HG003 | 14% | 10,650,000 | 45,166 | 133,600,000,000 | 50 |
| Nanopore ultra-long | HG004 | 13% | 7,640,251 | 48,518 | 69,900,000,000 | 26 |
| Pacbio Hifi | HG002 | 1% | 11,740,000 | 13,593 | 166,200,000,000 | 62 |
| Pacbio Hifi | HG003 | 1% | 8,425,983 | 14,977 | 129,500,000,000 | 48 |
| Pacbio Hifi | HG004 | 1% | 3,661,005 | 18,928 | 68,090,000,000 | 25 |

**Table S2:** PEPPER SNP calling statistics of HG002, HG003, and HG004.

| Sample | Homozygous SNP | Heterozygous SNP |
|---|---|---|
| HG002 ONT 60x | 1,784,079 | 2,262,494 |
| HG003 ONT 50x | 1,827,977 | 2,260,693 |
| HG004 ONT 25x | 2,848,376 | 2,358,752 |
| HG002 PB 60x | 1,886,251 | 2,366,423 |
| HG003 PB 50x | 1,883,702 | 2,349,164 |
| HG004 PB 25x | 2,003,887 | 2,427,183 |

**Table S3:** SNP-only phasing statistics of LongPhase, Margin, and WhatsHap.

| Sample | Program | SW | SW(%) | *Phased SNPs (%) | No. of Block | Block N50 (bp) | Max block (bp) | Process Time (min) |
|---|---|---|---|---|---|---|---|---|
| *HG002 ONT 60x* | WhatsHap | 134 | 0.15% | 99.89% | 534 | 13,353,904 | 47,091,786 | 840 |
| | Margin | 121 | 0.15% | 92.97% | 633 | 10,522,389 | 41,842,413 | 570 |
| | LongPhase | 141 | 0.16% | 99.07% | 334 | 20,581,870 | 93,223,336 | 22 |
| *HG003 ONT 50x* | WhatsHap | 99 | 0.13% | 99.87% | 761 | 9,932,828 | 32,822,369 | 764 |
| | Margin | 87 | 0.13% | 92.81% | 865 | 7,385,841 | 23,579,100 | 526 |
| | LongPhase | 99 | 0.13% | 98.88% | 550 | 12,160,204 | 47,152,303 | 20 |
| *HG004 ONT 25x* | WhatsHap | 87 | 0.12% | 99.78% | 567 | 12,534,188 | 43,182,284 | 580 |
| | Margin | 60 | 0.10% | 82.43% | 1391 | 3,792,651 | 15,767,031 | 263 |
| | LongPhase | 91 | 0.13% | 92.13% | 564 | 10,616,587 | 31,377,010 | 11 |
| *HG002 PB 60x* | WhatsHap | 147 | 0.16% | 99.86% | 13,770 | 421,184 | 2,692,512 | 550 |
| | Margin | 121 | 0.14% | 90.60% | 14,082 | 402,119 | 2,321,349 | 548 |
| | LongPhase | 149 | 0.16% | 99.88% | 13,666 | 425,621 | 2,692,512 | 22 |
| *HG003 PB 50x* | WhatsHap | 111 | 0.14% | 99.84% | 13,872 | 415,877 | 3,087,674 | 546 |
| | Margin | 98 | 0.13% | 93.94% | 13,998 | 402,993 | 3,087,674 | 493 |
| | LongPhase | 105 | 0.13% | 99.88% | 13,773 | 421,341 | 3,087,674 | 16 |
| *HG004 PB 25x* | WhatsHap | 99 | 0.12% | 99.81% | 11,852 | 479,746 | 3,303,232 | 436 |
| | Margin | 84 | 0.11% | 91.49% | 12,091 | 453,546 | 3,106,989 | 222 |
| | LongPhase | 100 | 0.12% | 99.87% | 11,755 | 487,657 | 3,303,232 | 10 |
| ***Phased SNPs(%) = Phased SNPs / total hetero SNP** | | | | | | | | |

**Table S4:** Statistics of SVs called by Sniffles and CuteSV on HG002, HG003, and HG004.

| | Sniffles | | CuteSV | |
|---|---|---|---|---|
| Sample | Homozygous SV | Heterozygous SV | Homozygous SV | Heterozygous SV |
| HG002 ONT 60x | 24,942 | 17,708 | 25,424 | 25,813 |
| HG003 ONT 50x | 21,939 | 16,921 | 21,986 | 24,251 |
| HG004 ONT 25x | 13,912 | 10,683 | 16,890 | 14,902 |
| HG002 PB 60x | 20,816 | 14,058 | 17,034 | 20,027 |
| HG003 PB 50x | 17,808 | 14,243 | 15,964 | 18,872 |
| HG004 PB 25x | 12,227 | 7,787 | 12,795 | 11,677 |

**Table S5:** Types of heterozygous SVs called by Sniffles.

| | DEL | INS | DUP | INV | DEL/INV | DUP/INS | INV/DUP | sum |
|---|---|---|---|---|---|---|---|---|
| HG002 ONT 60x | 10,143 | 7,280 | 60 | 36 | 1 | | | 17,708 |
| HG003 ONT 50x | 9,602 | 6,990 | 71 | 50 | 1 | | | 16,921 |
| HG004 ONT 25x | 6,317 | 4,168 | 40 | 29 | 1 | | | 10,683 |
| HG002 PB 60x | 6,034 | 7,193 | 77 | 58 | 19 | | | 14,058 |
| HG003 PB 50x | 6,183 | 7,260 | 79 | 68 | 22 | 1 | 1 | 14,243 |
| HG004 PB 25x | 3,270 | 3,987 | 40 | 43 | 29 | | | 7,787 |

**Table S6:** Statistics of SNP/SV co-phasing by LongPhase. The SVs called by Sniffles and CuteSVs are separately co-phased by LongPhase.

| | Sample | *Phased SVs (%) | No. of Block | Block N50 (bp) | Max block (bp) |
|---|---|---|---|---|---|
| PEPPER SNPs | HG002 ONT 60x | - | 334 | 20,581,870 | 93,223,336 |
| | HG003 ONT 50x | - | 550 | 12,160,204 | 47,152,303 |
| | HG004 ONT 25x | - | 564 | 10,616,587 | 31,377,010 |
| | HG002 PB 60x | - | 13,666 | 4,25,621 | 2,692,512 |
| | HG003 PB 50x | - | 13,773 | 4,21,341 | 3,087,674 |
| | HG004 PB 25x | - | 11,755 | 4,87,657 | 3,303,232 |
| PEPPER SNPs + Sniffles SVs | HG002 ONT 60x | 95.06% | 297 | 26,250,786 | 93,223,336 |
| | HG003 ONT 50x | 94.12% | 495 | 14,211,717 | 47,366,189 |
| | HG004 ONT 25x | 90.98% | 517 | 12,036,007 | 31,380,743 |
| | HG002 PB 60x | 90.55% | 13,438 | 431,824 | 2,692,512 |
| | HG003 PB 50x | 90.62% | 13,515 | 427,953 | 3,087,674 |
| | HG004 PB 25x | 89.38% | 11,620 | 495,549 | 3,303,232 |
| PEPPER SNPs + CuteSV SVs | HG002 ONT 60x | 94.75% | 287 | 26,250,786 | 93,223,336 |
| | HG003 ONT 50x | 94.99% | 482 | 14,519,944 | 47,152,303 |
| | HG004 ONT 25x | 92.95% | 525 | 11,479,601 | 31,385,793 |
| | HG002 PB 60x | 96.30% | 13,359 | 434,138 | 2,692,512 |
| | HG003 PB 50x | 96.31% | 13,503 | 428,346 | 3,087,674 |
| | HG004 PB 25x | 95.47% | 11,585 | 495,902 | 3,303,232 |
| ***Phased SVs (%) = Phased SVs / total hetero SV** | | | | | |

**Table S7:** Types of heterozygous SVs called by CuteSV.

| | DEL | INS | DUP | INV | TLC | sum |
|---|---|---|---|---|---|---|
| HG002 ONT 60x | 15,736 | 9,521 | 388 | 73 | 95 | 25,813 |
| HG003 ONT 50x | 14,285 | 9,435 | 364 | 72 | 95 | 24,251 |
| HG004 ONT 25x | 8,705 | 5,856 | 226 | 39 | 76 | 14,902 |
| HG002 PB 60x | 9,419 | 9,949 | 306 | 76 | 277 | 20,027 |
| HG003 PB 50x | 8,938 | 9,342 | 262 | 66 | 264 | 18,872 |
| HG004 PB 25x | 5,731 | 5,528 | 129 | 49 | 240 | 11,677 |
